# Lateralized vagal oxytocin signaling separately controls feeding and socioemotional functions via hypothalamic oxytocin signaling

**DOI:** 10.64898/2025.12.16.694527

**Authors:** Kengo Iba, Yuta Masuda, Rika Kitano, Kento Ohbayashi, Yukio Ago, Hirotaka Nagai, Tomoyuki Furuyashiki, Chikara Abe, Yusaku Iwasaki

## Abstract

Oxytocin neurons in the paraventricular hypothalamus (PVH^Oxt^) regulate feeding, anxiety, and social behaviors. Activation of Oxt receptor (Oxtr) -expressing vagal sensory neurons engages these PVH^Oxt^ neurons and improves hyperphagic obesity; however, their roles in anxiety and sociability remain unclear. Here, we activated vagal Oxtr-expressing neurons in male mice using a single intraperitoneal (IP) Oxt injection or chemogenetics. IP Oxt reduced anxiety-like behavior, enhanced social interaction, and suppressed feeding while activating both vagal sensory neurons and PVH^Oxt^ neurons. These effects were abolished by chemogenetic inhibition of PVH^Oxt^ neurons or central Oxtr blockade. Subdiaphragmatic vagotomy revealed lateralized functions: right-side vagotomy eliminated anxiolytic and prosocial effects, whereas left-side vagotomy blocked feeding suppression. Consistently, chemogenetic activation of left-sided neurons suppressed feeding, while right-sided activation reduced anxiety and increased sociability. These findings identify Oxtr-expressing vagal sensory neurons as a major peripheral pathway in which left- and right-sided inputs differentially control feeding and socioemotional behaviors.

## INTRODUCTION

Oxytocin (Oxt) is a peptide hormone composed of nine amino acids, produced primarily in the paraventricular nucleus (PVH) and supraoptic nucleus (SON) of the hypothalamus. A subset of Oxt-producing neurons projects to the posterior pituitary and releases Oxt into the circulation, where it acts as a classical neurohypophysial hormone to regulate peripheral physiological functions such as uterine contraction and milk ejection ^1^. In addition, many Oxt neurons send axons to various brain regions ^2^, where Oxt acts as a neurotransmitter or neuromodulator to regulate central functions. Centrally released Oxt contributes not only to the control of energy homeostasis including feeding behavior, glucose and lipid metabolism, and energy expenditure ^3^ but also to the modulation of mental functions such as anxiety and social behavior ^4^.

Vagal sensory nerves detect diverse signals arising from peripheral organs and convey this information to the central nervous system, thereby contributing to the regulation of feeding, metabolism, and mental functions ^5^. Recent single-cell transcriptomic analyses have revealed that vagal sensory neurons can be classified into multiple subclasses based on their gene expression profiles ^6,7^, providing a molecular basis for their functional diversity. Among these, oxytocin receptor (Oxtr)-expressing vagal sensory neurons have been shown to play a key role in the regulation of feeding ^6^.

We previously demonstrated that peripheral administration of Oxt activates Oxtr-expressing vagal sensory neurons, which in turn stimulate PVH^Oxt^ neurons, leading to improvements in hyperphagia, obesity, and type 2 diabetes ^8–10^. PVH^Oxt^ neurons are known to regulate not only feeding but also anxiety- and social behavior–related circuits ^11,12^. Conversely, obesity and type 2 diabetes are established risk factors for depression ^13–15^. Thus, activation of PVH^Oxt^ neurons via Oxtr-expressing vagal sensory neurons may not only ameliorate metabolic disorders but also improve anxiety- and social-behavior deficits associated with depression.

In this study, we investigated whether peripheral Oxt administration, through activation of vagal sensory inputs to PVH^Oxt^ neurons, influences not only feeding but also anxiety and social behaviors. Because several physiological functions are reported to be asymmetrically regulated by the left or right vagal sensory branch ^16–18^, we further examined the lateralized functions of Oxtr-expressing vagal sensory neurons. We demonstrate that left Oxtr-expressing vagal afferents mediate feeding suppression, whereas the right counterparts regulate anxiety and social behaviors, revealing functional lateralization of Oxtr-expressing vagal sensory pathways.

## RESULTS

### Single intraperitoneal Oxt injection reduces anxiety, enhances social behavior, and suppresses food intake via peripheral Oxt receptors

To examine the effects of peripherally administered Oxt on anxiety- and social-related behaviors, normal C57BL/6J male mice received a single intraperitoneal (IP) injection of Oxt, followed 1 h later by either the elevated plus maze (EPM) or the three-chamber social interaction test (SIT). Oxt (400 nmol/kg) significantly increased the time spent in the open arms, an index of anxiolytic behavior, in the EPM (Figures 1A and 1B). In contrast, no statistically significant changes were observed in the time spent in the closed arms, an index of anxiety, or in total distance traveled, an index of locomotor activity (Figures 1C and 1D). The Oxt-induced anxiolytic response was abolished by co-administration of the peripherally acting Oxt receptor (Oxtr) antagonist L-371,257 at 20 µmol/kg (Figures 1E‒G), whose penetration across the blood–brain barrier is limited ^19,20^. In the SIT (Figure 1H and S1), Oxt IP significantly increased the time spent nose-poking in the sniffing zone adjacent to target cage containing a mouse (Figure 1I, mouse side [M] in saline vs. Oxt), without affecting total distance traveled (Figure 1J), thereby indicating enhanced prosocial behavior. This Oxt-induced prosocial response was also abolished by co-administration of L-371,257 (Figures 1K and 1L).

**Figure 1.**
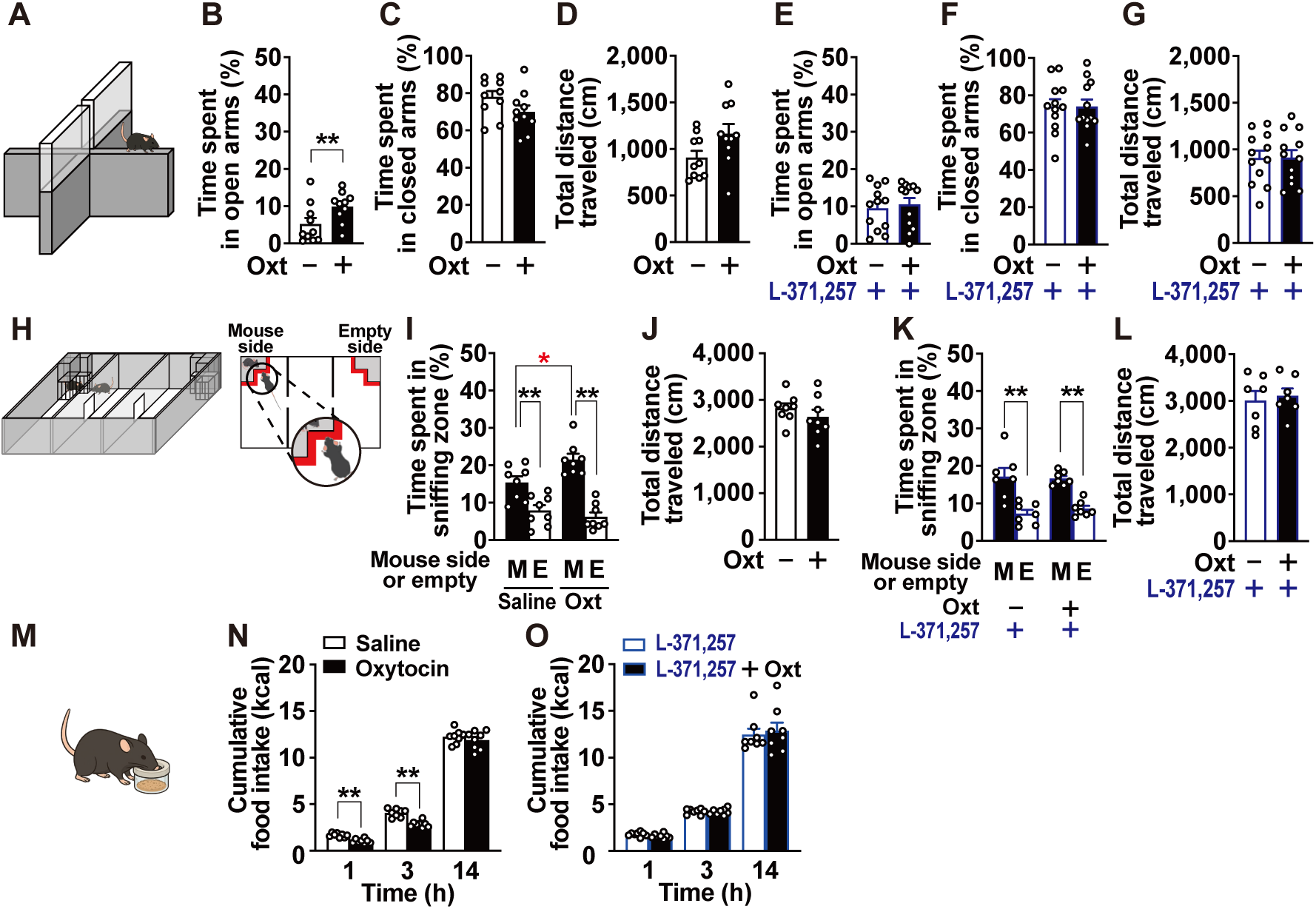
A single intraperitoneal injection of oxytocin reduces anxiety-like behavior, enhances social interaction, and suppresses food intake via peripheral oxytocin receptor signaling in normal C57BL/6J male mice. (**A–D**) Effect of oxytocin (Oxt) on anxiety-like behavior in elevated plus maze (EPM). (**A**) Schematic representation of EPM. (**B–D**) Time spent in open arms, time spent in closed arms, and total distance traveled during a 5-min test conducted 1 h after intraperitoneal (IP) injection of Oxt (400 nmol/kg) or saline. In the figures, “Oxt, –” indicates IP injection of saline. n = 10. **p < 0.01 by unpaired t-test (B). (**E–G**) Inhibitory effect of Oxt receptor antagonist L-371,257 on Oxt’s effect in EPM. Saline-based solutions containing 20% DMSO, together with either L-371,257 (10 mg/kg) and Oxt (400 nmol/kg) or L-371,257 alone, were administered intraperitoneally. n = 12. (**H–L**) Effect of Oxt on social behavior in three-chamber social interaction test (SIT). (**H**) Schematic representation of SIT. (**I and J**) Time spent nose-poking in the sniffing zone and total distance traveled during a 10-min test performed 1 h after IP injection of Oxt (400 nmol/kg) or saline. M and E indicate mouse side or empty side, respectively. n = 8. *p < 0.05, **p < 0.01 by one-way ANOVA followed by Tukey’s test (I). (**K and L**) Effect of L-371,257 (10 mg/kg, IP) co-administration on Oxt-induced social interaction. n = 12. **p < 0.01 by one-way ANOVA followed by Tukey’s test (K). (**M–O**) Effect of Oxt on food intake during dark period in mice fasted for 3 hours. (**M**) Schematic of feeding test. (**N**) Cumulative food intake during 1–14 h immediately after IP injection of Oxt (400 nmol/kg) or saline. n = 8. **p < 0.01 by unpaired t-test (N). (**O**) Effect of L-371,257 (10 mg/kg, IP) co-administration on the Oxt-induced anorexigenic effect. n = 8.

A single IP injection of Oxt reportedly suppressed short-term food intake ^9^. When administered at the onset of the dark phase, Oxt (400 nmol/kg) significantly reduced cumulative food intake at 1 and 3 h after IP injection (Figures 1M and 1N). This anorexigenic effect was completely abolished by co-administration of L-371,257 (Figure 1O). These results demonstrate that a single IP injection of Oxt exerts anxiolytic, prosocial, and anorexigenic effects, all mediated through peripheral Oxtr signaling.

### IP Oxt activates vagal sensory neurons, the nucleus tractus solitarius, and oxytocin neurons in the paraventricular nucleus of the hypothalamus

To determine whether a single IP injection of Oxt activates vagal sensory neurons *in vivo*, we performed histological analyses using pERK1/2 as a marker of neuronal activity ^21^. A single IP injection of Oxt (400 nmol/kg) significantly increased the proportion and number of pERK1/2-positive neurons in the left nodose ganglion (NG; Figures 2A–2B, 2D), right NG (Figures 2E–2F, 2H), and medial nucleus tractus solitarius (medial NTS) (Figures 2I–2J, 2L) 15 min after injection. There was no difference between the left and right NTS in the increase of c-Fos–positive neurons induced by IP Oxt administration. This Oxt-induced activation was completely abolished by co-administration of the Oxtr antagonist L-371,257 (Figures 2C–2D, 2G–2H, 2K–2L).

**Figure 2.**
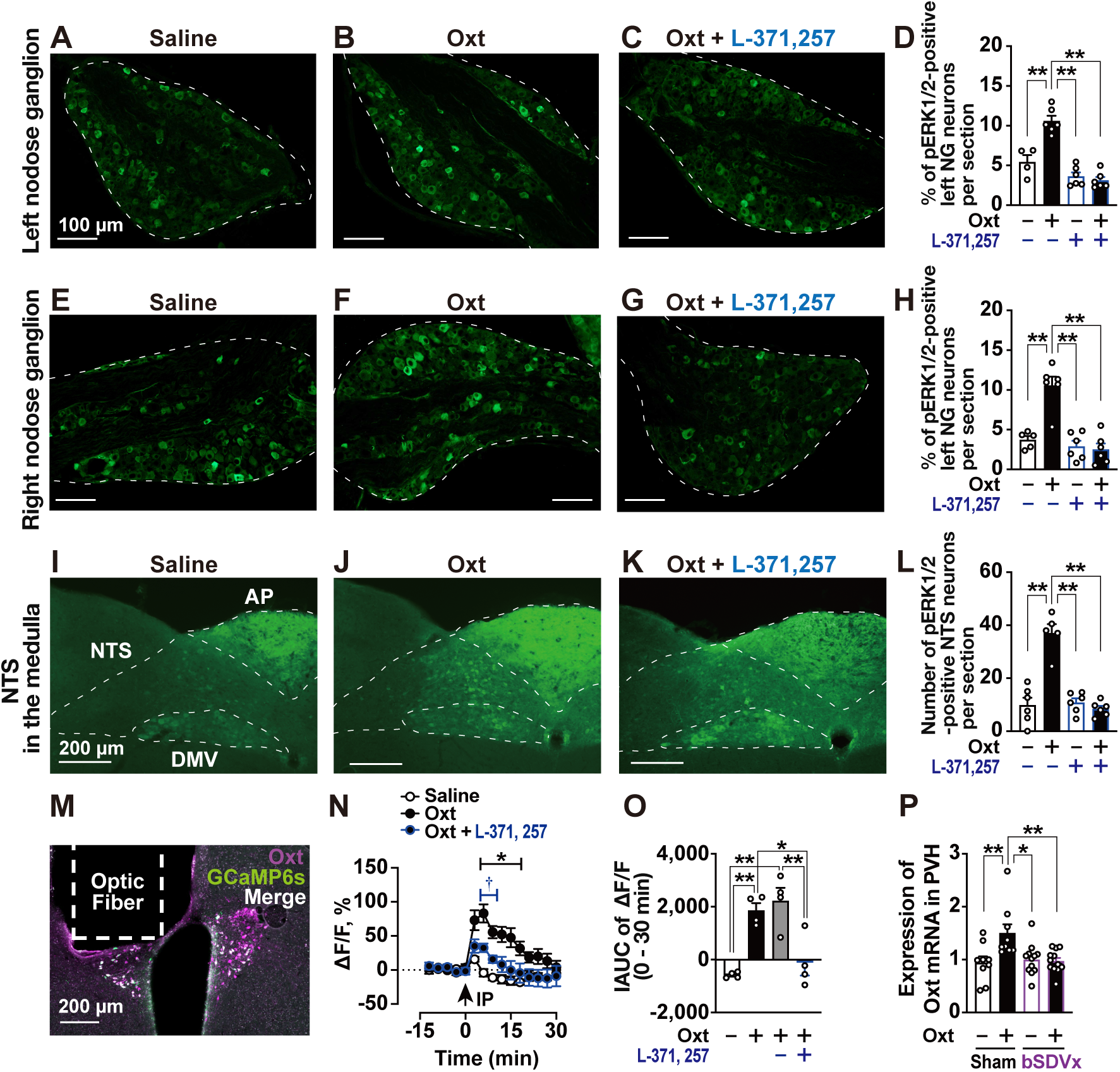
IP injection of Oxt activates nodose ganglia (NGs), nucleus tractus solitarius (NTS), and Oxt neurons in paraventricular nucleus of the hypothalamus (PVH^Oxt^ neurons) via peripheral Oxt receptor signaling. (**A–L**) Histological analysis of neuronal activation induced by Oxt IP injeciton using pERK1/2 as a marker of neural activation. Representative images of pERK1/2 immunostaining in left NG (**A–C**), right NG (**E–G**), and bilateral medial NTS (**I–K**) 15 min after IP injection of saline, Oxt (400 nmo/kg), or Oxt combined with Oxt receptor antagonist L-371,257 (10 mg/kg) in C57BL/6J male mice. Scale bars: 100 µm in NGs and 200 µm in NTS. AP, area postrema; DMV, dorsal motor nucleus of the vagus. (**D, H and L**) Percentage of pERK1/2-positive neurons in left NG (**D**) and right NG (**H**). Number of pERK1/2-positive neurons in NTS (**L**). Because L-371,257 was prepared in a DMSO solution, the injected solutions indicated as “L-371,257, –” and “L-371,257, +” contained 20% DMSO. n = 4–6. **p < 0.01 by one-way ANOVA followed by Tukey’s test (D, H and L). (**M–O**) Fiber photometry recordings of PVH^Oxt^ neuronal activity. (**M**) Representative coronal section of the PVH showing location of optical fiber and expression of GCaMP6s (green) in PVH^Oxt^ neurons (magenta). Scale bars: 200 µm. (**N**) Time course of photometry signals (ΔF/F) recorded every 3 min from 15 min before to 30 min after IP injection of saline or Oxt (400 nmol/kg), with or without L-371,257 (10 mg/kg). n = 4. *p < 0.05 (Oxt vs Saline) and †p < 0.05 (Oxt vs Oxt + L-371,257) by two-way ANOVA followed by Dunnett’s test (N). (**O**) Incremental area under the curve (IAUC) of ΔF/F signals. The gray bar “L-371,257, –” indicates Oxt dissolved in the vehicle containing 20% DMSO, which is shown in (O) but not included in (N). n = 4. *p < 0.05, **p < 0.01 by one-way ANOVA followed by Tukey’s test (O). (**P**) Relative *Oxt* mRNA expression in PVH 2 h after IP injection of Oxt (400 nmol/kg) or saline in sham-operated and bilateral subdiaphragmatic vagotomized (bSDVx) mice. 36b4 was used as an internal control. n = 8–12. *p < 0.05, **p < 0.01 by one-way ANOVA followed by Tukey’s test (P).

We previously reported that IP injection of Oxt activates PVH^Oxt^ neurons via vagal sensory pathways, as assessed by c-Fos immunostaining ^10^. In the present study, we evaluated neuronal activity of Oxt neurons in the paraventricular nucleus of the hypothalamus (PVH^Oxt^ neurons) in real time using *in vivo* fiber photometry in freely moving mice that received a unilateral PVH-targeted injection of AAV9 expressing GCaMP6s under the Oxt promoter ^22^ (Figure 2M). IP injection of Oxt transiently increased fluorescence intensity, reflecting cytosolic Ca^2+^ signals in PVH^Oxt^ neurons, with a peak at ∼6 min and a gradual return to baseline within 30 min (Figure 2N). Compared with saline, Oxt induced a significantly greater response, as confirmed by incremental area under the curve (IAUC) analysis (Figure 2O). This Ca^2+^ response was completely suppressed by co-administration of L-371,257 (Figures 2N and 2O).

Furthermore, 2 h after IP injection, Oxt significantly increased *Oxt* mRNA expression in the PVH but not in the supraoptic nucleus (Figure S2). This Oxt-induced upregulation of PVH *Oxt* mRNA was abolished by bilateral subdiaphragmatic vagotomy (Figure 2P). Together, these findings demonstrate that IP injection of Oxt rapidly activates PVH^Oxt^ neurons through the activation of Oxtr-expressing vagal afferents.

### PVH^Oxt^ neurons and central Oxtr signaling are essential for Oxt-induced anxiolytic and prosocial responses

To examine whether PVH^Oxt^ neurons contribute to Oxt-induced anxiolytic and prosocial responses, we used *Oxt-ires-Cre* mice bilaterally injected with AAV2-hSyn-DIO-hM4Di-mCherry in the PVH to selectively inhibit these neurons using an inhibitory DREADD (Figure 3A). First, we validated that the inhibitory DREADD system effectively suppressed Oxt-induced activation of PVH^Oxt^ neurons, as assessed by c-Fos expression. In wild-type mice without AAV injection, pretreatment with hM4Di agonist clozapine-*N*-oxide (CNO, 1 mg/kg, IP) followed by IP injection of Oxt at 400 nmol/kg significantly increased c-Fos expression in PVH^Oxt^ neurons (Figures 3B and 3C). In contrast, in *Oxt-ires-Cre* mice expressing hM4Di in PVH^Oxt^ neurons (PVH^Oxt^-Gi mice), pretreatment with CNO completely abolished Oxt-induced c-Fos expression (Figures 3B and 3C). These results indicate that the DREADD system effectively inhibited PVH^Oxt^ neuronal activity. Next, when PVH^Oxt^-Gi mice were pretreated with vehicle followed by a single IP injection of Oxt (400 nmol/kg), food intake was suppressed (Figure 3D), anxiolytic behavior (Figure 3E) and social behavior (Figure 3F) were significantly increased. These effects were markedly attenuated by pretreatment with CNO (Figures 3D–F). These results show that PVH^Oxt^ neurons are required for Oxt-induced anorexigenic, anxiolytic, and prosocial effects.

**Figure 3.**
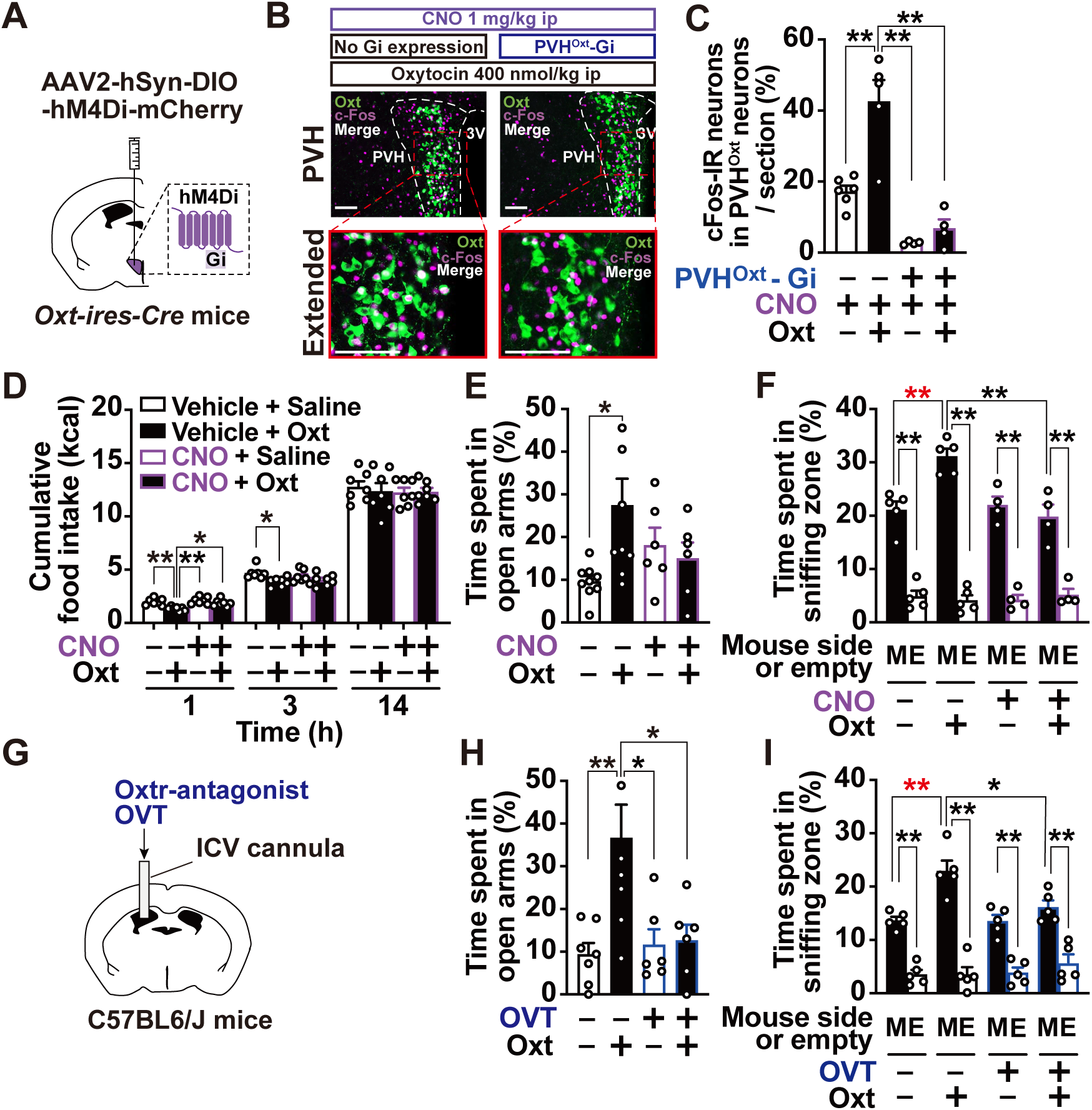
Anxiolytic, prosocial, and anorexigenic effects of single IP Oxt injection via PVH^Oxt^ neurons and central Oxt receptors. (**A–F**) Effects of chemogenetic inhibition of PVH^Oxt^ neurons on Oxt-induced anxiolytic, prosocial, or anorexigenic responses. (**A**) Experimental scheme for chemogenetic inhibition using bilateral hM4Di-mCherry AAV injection into PVH of *Oxt-ires-Cre* mice. (**B**) Representative images of the PVH showing double immunostaining for c-Fos (magenta) and Oxt neurons (green) in mice with or without hM4Di expression in PVH^Oxt^ neurons. Mice were injected IP with clozapine-*N*-oxide (CNO, 1 mg/kg), followed 1 h later by either saline or Oxt (400 nmol/kg), and brains were collected 90 min after the second injection. (**C**) Percentage of c-Fos-positive neurons in PVH^Oxt^ neurons. PVH^Oxt^-Gi (–), wild-type C57BL/6J mice; PVH^Oxt^-Gi (+), *Oxt-ires-Cre* mice expressing hM4Di in PVH; Oxt (–), saline IP; Oxt (+), Oxt IP. (**D–F**) Effects of pretreatment with CNO (3 mg/kg) on Oxt-induced anorexigenic (**D**), with CNO (1 mg/kg) on Oxt-induced anxiolytic (**E**), or prosocial (**F**) responses in *Oxt-ires-Cre* mice expressing hM4Di in PVH^Oxt^ neurons. (**D**) Cumulative food intake immediately after IP injection of Oxt (400 nmol/kg) or saline. n = 7. (**E and F**) Time spent in open arms in EPM (**E,** n = 6 or 9) and time spent nose-poking in sniffing zone in SIT (**F**, n = 4–5) 1 h after IP injection of Oxt (400 nmol/kg) or saline. (**G–I**) Effects of intracerebroventricular (ICV) injection of saline (2 μL/mouse) or Oxt receptor antagonist OVT ((d(CH_2_)_5_^1^, Tyr(Me)^2^, Orn^8^)-oxytocin, 3.7 nmol/2 μL/mouse) on Oxt-induced anxiolytic or prosocial responses in C57BL/6J mice. ICV injection was carried out 15 min prior to Oxt IP injection, and behavioral tests were conducted 1 h after Oxt administration. (**G**) Schematic of unilateral ICV cannulation into lateral ventricle. (**H and I**) Time spent in open arms in EPM (**H,** n = 6–8) and time spent nose-poking in sniffing zone in SIT (**I**, n = 5). In figure D–F, “CNO, –” indicates vehicle solution containing 1% DMSO. In figure H–I, “OVT, –” indicates saline ICV injection. In figure F and I, “M” and “E” indicate mouse side or empty side, respectively. *p < 0.05, **p < 0.01 by one-way ANOVA followed by Tukey’s test (C–F, H and I).

To assess the contribution of central Oxtr signaling, we administered the Oxtr antagonist OVT, (d(CH_2_)_5_^1^,Tyr(Me)^2^,Orn^8^)-oxytocin, intracerebroventricularly (ICV) via cannula implanted into the lateral ventricle (Figure 3G). The Oxt-induced increase in open-arm time in the EPM and enhancement of social interaction in the SIT were completely abolished by ICV pretreatment with OVT (3.7 nmol/mouse, Figures 3H and 3I). We previously reported that the anorexigenic effect of Oxt is also blocked by ICV OVT ^10^. These findings indicate that peripheral Oxt administration regulates feeding, anxiety, and social behaviors through both peripheral and central Oxtr signaling.

### Peripheral Oxt actions are asymmetrically mediated by left and right vagal afferents

We next examined the involvement of subdiaphragmatic vagal nerves in the anxiolytic, prosocial, and anorexigenic effects of peripheral Oxt using mice subjected to left- or right-sided subdiaphragmatic vagotomy. In sham-operated mice (Figure 4A), IP Oxt significantly increased time spent in the open arms of the EPM (Figure 4B), increased the time spent nose-poking in the sniffing zone adjacent to the target cage [M] in the SIT (Figure 4C), and suppressed cumulative food intake (Figure 4D). In left-sided subdiaphragmatic vagotomized mice (Left-SDVx, Figure 4E), the anxiolytic (Figure 4F) and prosocial (Figure 4G) effects of Oxt were preserved, whereas its anorexigenic effect was abolished (Figure 4H). In contrast, in right-sided subdiaphragmatic vagotomized mice (Right-SDVx, Figure 4I), the anxiolytic (Figure 4J) and prosocial (Figure 4K) effects of Oxt were abolished, while the anorexigenic effect was maintained (Figure 4L). These results demonstrate that peripheral Oxt suppresses feeding via the left vagus nerve, whereas it reduces anxiety and enhances social behavior via the right vagus nerve.

**Figure 4.**
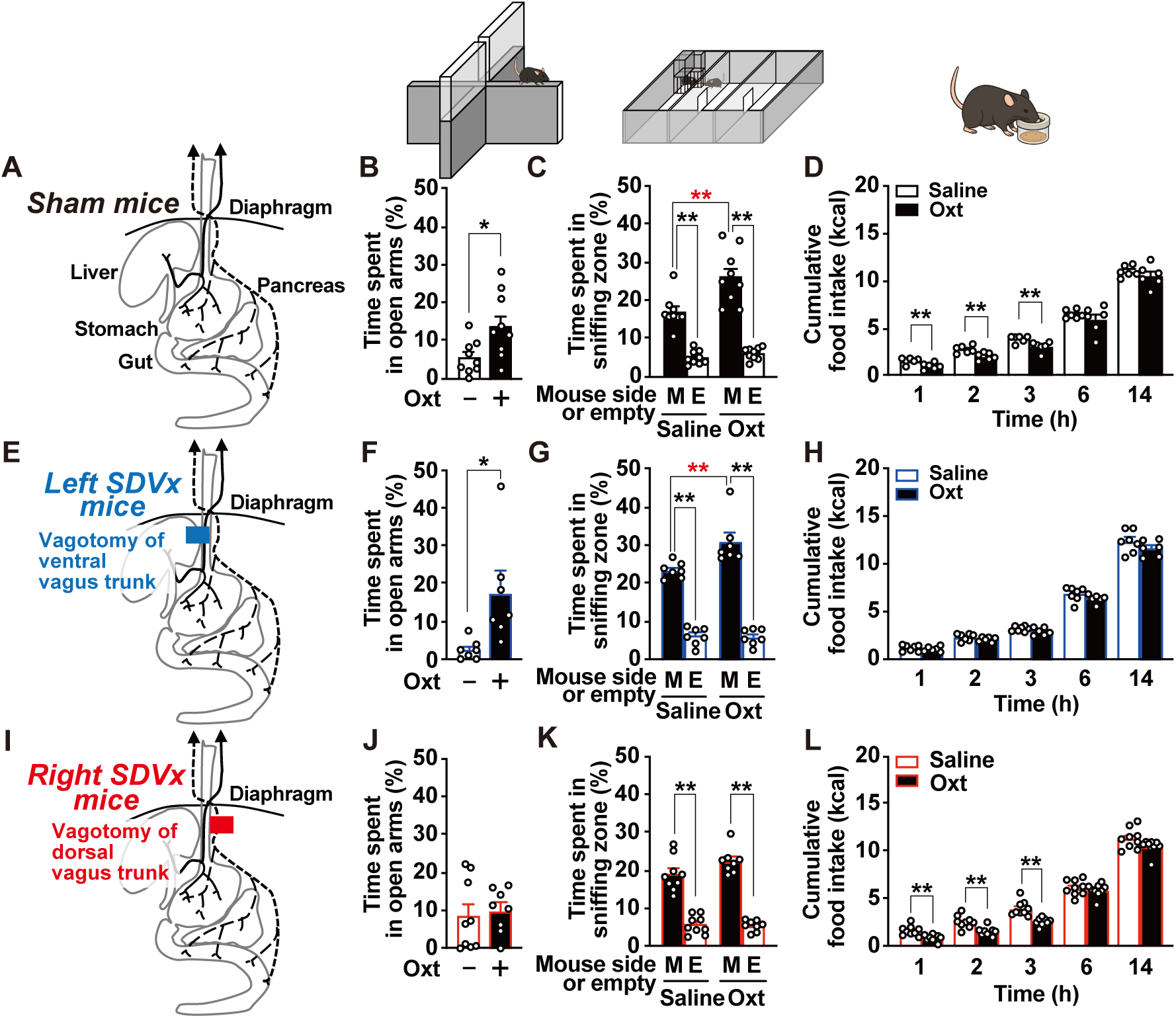
Anxiolytic and prosocial effects of IP Oxt are mediated by right vagal afferents, whereas its anorexigenic effect is mediated by left vagal afferents. (**A, E, and I**) Schematic diagram of the subdiaphragmatic vagal nerve and the site of transection. Sham operation (**A–D**), left-side (**E–H,** Left-SDVx) or right-side (**I–J,** Right-SDVx) subdiaphragmatic vagotomy was performed in C57BL/6J male mice. (**B, F, and J**) Oxt-induced anxiolytic effect in EPM, shown as time spent in open arms 1 h after Oxt IP (400 nmol/kg) in sham (n = 9), Left-SDVx (n =7), or Right-SDVx (n = 8–9) mice. (**C, G, and K**) Oxt-induced prosocial effect in SIT, shown as time spent nose-poking in sniffing zone 1 h after Oxt IP (400 nmol/kg) in sham (n = 9), Left-SDVx (n =7), or Right-SDVx (n = 8–9) mice. M and E indicate mouse side or empty side, respectively. (**D, H, and L**) Oxt-induced anorexigenic effect, shown as cumulative food intake immediately after Oxt IP (400 nmol/kg) in sham (n = 6), Left-SDVx (n =7), or Right-SDVx (n = 9) mice. *p < 0.05, **p < 0.01 by unpaired t-test (B, D, F and L). *p < 0.05, **p < 0.01 by one-way ANOVA followed by Tukey’s test (C, G and K).

### Selective activation of left and right Oxtr-expressing vagal afferent subclasses drives distinct behavioral responses

Finally, we examined the functions regulated by left versus right Oxtr-expressing vagal afferents. An AAV vector encoding Cre-dependent excitatory DREADD (hM3Dq) was locally injected into either the left or right nodose NG of *Oxtr-iCre* mice (Figures 5A and 5B). Beginning 3 weeks after injection, when hM3Dq expression was stable (Figures 5C–5F), these neurons were selectively activated by IP administration of CNO at 1 mg/kg, followed by analysis of anxiety-like, social, and feeding behaviors.

**Figure 5.**
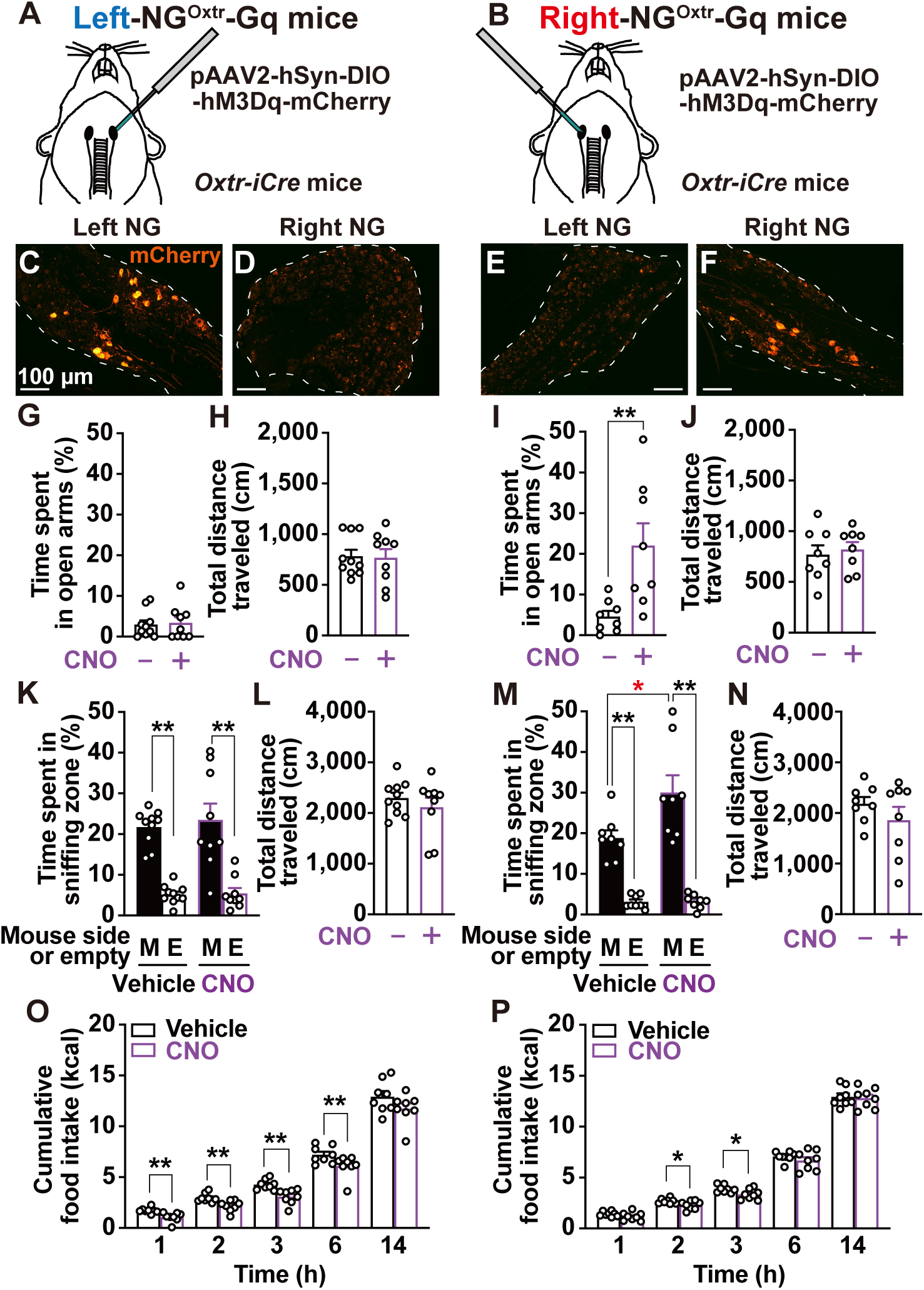
Chemogenetic activation of Oxt receptor–expressing vagal afferents reveals lateralized regulation of distinct functions. (**A, B**) Schematic of AAV microinjection expressing Cre-dependent expression of hM3Dq and mCherry into the unilateral left (**A**, Left-NG^Oxtr^-Gq mice) or right (**B**, Right-NG^Oxtr^-Gq mice) NG of *Oxt receptor-iCre* (*Oxtr-iCre*) mice. (**C–F**) Representative images of mCherry expression (red) in left NG (**C, E**) or right NG (**D, F**) after AAV injection, indicating that robust mCherry expression was observed only in the injected site (**C, F**). Scale bars, 100 µm. (**G–J**) Assessment of anxiety-like behavior using EPM. Time spent in the open arms (**G and I**) and total distance traveled (**H and J**) 1 h after IP injection of CNO (1 mg/kg) or vehicle in Left-NG^Oxtr^-Gq mice (n = 9–10) or Right-NG^Oxtr^-Gq mice (n = 8). **p < 0.01 by unpaired t-test (I). (**K–N**) Social interaction test (SIT) 1 h after IP injection of CNO (1 mg/kg) or vehicle in Left- (n = 9–10) and Right-NG^Oxtr^-Gq mice (n = 8). In figures *K* and *M*, M and E indicate mouse side or empty side, respectively. *p < 0.05, **p < 0.01 by one-way ANOVA followed by Tukey’s test (K and M). (**O and P**) Cumulative food intake immediately after CNO IP (1 mg/kg) in Left- (n = 8) and Right-NG^Oxtr^-Gq mice (n = 8). *p < 0.05, **p < 0.01 by unpaired t-test (O and P).

Activation of left Oxtr-expressing vagal afferents did not affect behavioral measures in the EPM (Figures 5G and 5H) or SIT (Figures 5K and 5L), but significantly reduced cumulative food intake for 1–6 h after CNO administration (Figure 5O). In contrast, activation of right Oxtr-expressing vagal afferents significantly increased open-arm time in the EPM (Figure 5I) and sniffing time toward the target mouse in the SIT (Figure 5M), without affecting total distance traveled in either test (Figures 5J and 5N). Right-sided activation also reduced cumulative food intake, but only at 2–3 h after CNO injection, and this effect was limited compared with that induced by left-sided activation (Figures 5P vs. 5O). Therefore, these findings provide direct evidence that Oxtr-expressing vagal afferents are divided into lateralized subclasses, with left-sided afferents primarily mediating anorexigenic effects and right-sided afferents mediating anxiolytic and prosocial effects.

## DISCUSSION

In this study, we demonstrate that a single IP administration of Oxt not only suppresses food intake but also reduces anxiety-like behavior and enhances sociability. These effects require activation of vagal sensory neurons expressing Oxtr, which in turn drives PVH^Oxt^ neuronal activity to regulate feeding, anxiety, and social behaviors. Importantly, by using DREADD technology to selectively manipulate the activity of Oxtr-expressing vagal afferents on each side, we identify a functional lateralization: left-sided neurons mediate feeding suppression, whereas right-sided neurons control anxiety and sociability. Thus, peripheral Oxtr signaling engages not a single uniform pathway, but a lateralized vagal network that differentially orchestrates distinct behavioral outputs.

We previously reported that vagal sensory neurons expressing Oxtr play a key role in suppressing food intake and preventing hyperphagia ^9^, and that this feeding control critically depends on PVH^Oxt^ neurons ^10^. In the present study, we show that stimulation of Oxtr-expressing vagal afferents reduces anxiety and enhances sociability through the activation of PVH^Oxt^ neurons. Recent single-cell RNA-sequencing analyses of nodose ganglia have identified distinct subclasses of Oxtr-expressing neurons ^6,7^. Several studies have demonstrated that chemogenetic activation of these Oxtr-expressing neurons suppresses feeding ^6,23^. Notably, Bai *et al.* further showed that activation of these neurons rapidly inhibits the activity of neuropeptide Y (NPY)/agouti-related peptide (AgRP) neurons in the arcuate nucleus of the hypothalamus (ARC), the principal orexigenic neurons that exert the strongest influence on feeding behavior. PVH^Oxt^ neurons reportedly activate ARC^POMC^ neurons ^24^, whereas no direct evidence indicates that PVH^Oxt^ neurons control AgRP neurons. Thus, Oxtr-expressing vagal afferents may influence AgRP neurons indirectly through PVH^Oxt^-related pathways involving ARC^POMC^ neurons.

Scott *et al.* reported that bilateral chemogenetic activation of Oxtr-positive vagal afferents increases anxiety, presumably through activation of corticotropin-releasing hormone (CRH) neurons in PVH ^23^. In contrast, our unilateral activation experiments revealed that stimulation of right-sided Oxtr-expressing vagal afferents decreased anxiety as shown in Figure 5. Thus, the opposing outcomes may reflect differential behavioral consequences of bilateral versus unilateral activation of Oxtr-expressing vagal afferents. Moreover, we found that IP Oxt injection reduced anxiety and enhanced sociability through vagal activation as demonstrated in Figure 4. This IP Oxt treatment increased *Oxt*, but not *Crh*, mRNA levels in the PVH (Figure S2A). Because peripheral Oxt has an extremely short half-life (1–2 min; ^25^), IP-administered Oxt likely acts preferentially on Oxtr-expressing vagal afferents innervating abdominal organs. Together, these findings suggest that activation of abdominal Oxtr-expressing vagal sensory neurons reduces anxiety and enhances social behavior.

IP injection of Oxt suppressed cumulative food intake for 3 h and reduced anxiety and enhanced social behavior at 1 h after injection. These effects were mediated by the activation of PVH^Oxt^ neurons. However, fiber photometry revealed that this activation lasted for only ∼15 min, indicating a clear temporal dissociation between PVH^Oxt^ neuronal activity and the emergence of behavioral and physiological effects. This dissociation suggests that PVH^Oxt^ activation engages downstream circuits ^2,26^ that operate on longer timescales to regulate several physiological functions. Future studies on the projection targets of PVH^Oxt^ neurons and the activity dynamics of these downstream regions will be essential to explain this temporal gap.

One of the key findings of this study is the clear functional lateralization within Oxtr-expressing vagal sensory neuron subclasses. Previous study has shown that vagal laterality contributes to distinct physiological functions: the left vagus regulates feeding behavior ^16^ and gut immune homeostasis through peripheral regulatory T cells ^17^, whereas the right vagus is involved in reward-related behavior ^18^. Our results extend this concept by demonstrating that left-sided Oxtr-expressing vagal afferents primarily regulate feeding, whereas right-sided counterparts control anxiety and social behaviors. Importantly, all of these lateralized functions were markedly attenuated when PVH^Oxt^ neurons were inhibited or when an Oxtr antagonist was intracerebroventricularly administered ^10^, highlighting a central role for the vagal–PVH^Oxt^ circuit in coordinating both feeding and socioemotional behaviors. Although we identified robust functional asymmetry at the level of vagal sensory inputs, no evidence currently supports lateralized functions of PVH^Oxt^ neurons themselves. PVH^Oxt^ neurons are thought to comprise functionally distinct subclasses, with projections to the arcuate nucleus, central amygdala, ventral tegmental area, and other targets, each contributing to the regulation of different physiological processes ^2,26^. Future studies are needed to determine whether higher-order hypothalamic circuits also exhibit functional asymmetry, and how such brain-level lateralization may contribute to the pathophysiology of hyperphagic obesity, metabolic disorders, and neuropsychiatric conditions including autism.

Intranasal Oxt has been reported to exert therapeutic effects on hyperphagia and obesity ^3,27,28^, as well as on neuropsychiatric disorders such as anxiety disorders ^29^, autism spectrum disorder and social anxiety disorder ^30,31^. In parallel, extensive animal studies have demonstrated that PVH^Oxt^ neurons are critical regulators of feeding behavior ^32^, body weight ^33^, anxiety-like behavior ^11^, and social behavior ^12^. Because the multiple therapeutic effects of intranasal Oxt parallel the diverse physiological functions regulated by PVH^Oxt^ neurons, PVH^Oxt^ neurons may contribute to the mechanisms underlying intranasal Oxt treatment. Intranasal Oxt reaches the cerebrospinal fluid in only a small amount, with a modest increase of approximately 1.5-fold, and may act directly on the brain, whereas a much larger proportion enters the systemic circulation ^34^. However, circulating Oxt has an extremely short half-life ^25^, and only ∼0.002% crosses the blood–brain barrier ^35^. Notably, Oxtr-expressing vagal sensory neurons are widely distributed in the periphery and influence PVH^Oxt^ neuron activity. Thus, intranasally administered Oxt might act through peripheral vagal afferent pathways to modulate PVH^Oxt^ neurons and thereby produce its therapeutic benefits.

Vagus nerve stimulation (VNS) is an established therapy for drug-resistant epilepsy ^36^, and recent reports indicate therapeutic benefits for treatment-resistant depression ^37^, anxiety disorders ^38^, and obesity ^39^. However, the underlying mechanisms remain largely unclear. Our finding that stimulation of Oxtr-expressing vagal sensory neurons regulates feeding, anxiety, and social behaviors through PVH^Oxt^ neurons may represent one mechanism contributing to the therapeutic effects of VNS. Clinically used VNS for epilepsy and depression predominantly targets the left cervical vagus nerve to avoid cardiovascular side effects ^40^. In contrast, our results reveal a clear functional lateralization within Oxtr-expressing vagal afferents: the left side primarily controls feeding behavior, whereas the right-side controls emotional and social behaviors. These findings suggest that future neuromodulation strategies targeting vagal afferents should consider this lateral specialization to achieve selective and minimally invasive therapeutic effects. Moreover, given the high comorbidity of obesity and depression ^15^, appropriately stimulating both sides of Oxtr-expressing vagal afferents may offer the possibility of treating multiple disorders simultaneously.

In this study, we show that activation of Oxtr-expressing vagal sensory neurons contributes not only to feeding suppression but also to anxiolytic and prosocial effects. These findings suggest that this Oxtr-expressing neuronal population plays multifaceted roles in the regulation of physiological and behavioral processes. However, little is known about the physiological conditions under which these neurons are activated. Oxt, the endogenous ligand for Oxtr, is expressed not only in the central nervous system but also in some sensory, enteric, and sympathetic neurons ^41,42^ ^43^, as well as in several peripheral organs such as the heart, testes, adrenal glands, and thymus ^44^. Understanding how Oxt production in these tissues is regulated and how it activates Oxtr-expressing vagal sensory neurons will be essential for elucidating the physiological significance of this pathway.

### Limitations of study

Several limitations should be noted. (1) The study was performed only in male mice, and potential sex differences remain to be addressed. (2) We focused on acute activation of Oxtr-expressing vagal sensory neurons by IP Oxt or DREADD technique, therefore the effects of chronic activation were not assessed. (3) Our findings are based on healthy mice, and the efficacy of activating Oxtr-expressing vagal sensory nerves in psychiatric disease models remains unknown. (4) The detailed function of the circuit linking vagal sensory neurons to the NTS, PVH^Oxt^ neurons, and downstream targets have yet to be defined. (5) Although we identified functional lateralization of vagal sensory nerves, how left and right vagal signals are integrated or segregated in the brain is still unclear.

## RESOURCE AVAILABILITY

### Lead contact

Further information and requests for resources, data, and code should be directed to and will be fulfilled by the lead contact, Yusaku Iwasaki.

### Materials availability

This study did not generate new, unique reagents.

### Data and code availability

This paper does not report original code. Any additional information required to reanalyze the data reported in this paper is available from the lead contact upon request.

## ACKNOWLEDGMENTS

The authors thank Drs. Hirokazu Hirai and Ayumu Konno (Gunma University) for developing the AAV9-OTp-GCaMP6s vector, and Dr. Kazunari Miyamichi (RIKEN Center for Biosystems Dynamics Research) for technical advice on fiber photometry and generously providing the AAV. We also thank all laboratory members (Kyoto Prefectural University) for their experimental support.

This study was supported in part by the Grant-in-Aid for Core Research for Evolutional Science and Technology (CREST, JPMJCR21P1 to Y.I.) from the Japan Science and Technology Agency (JST); by the Grant-in-Aid for JSPS Fellows (25KJ2028 to K.I.) from the Japan Society for the Promotion of Science (JSPS); by the Adaptable and Seamless Technology transfer Program through Target-driven R&D (A-STEP, JPMJTR20UT to Y.I.) from JST; by the Japan Agency for Medical Research and Development (AMED, JP24ym0126815 to Y.I., 25zf0127010, 25zf0127012, 25wm0625324 to T.F.); by the Grant-in-Aid for Scientific Research (24K22086 to T.F.) from JSPS; Moonshot R&D (JPMJMS239F to T.F.) from JST; by a Daiichi Sankyo Foundation of Life Science to Y.I..

## AUTHOR CONTRIBUTIONS

K.I., Y.M. and Y.I. developed concept and designed the study. K.I., Y.M., R.K., K.O. and Y.I. performed experiments and analyzed data. K.I. Y.A., H.N., T.F. and Y.I. optimized the behavioral pharmacological experimental methods in this study. K.O. C.A., and Y.I. contributed to establishing the microinjection method into the nodose ganglion in mice. K.I., Y.M. and Y.I. prepared figures, interpreted the results of the experiments, and drafted the manuscript. All of the authors edited the final draft. All authors have read and agreed to the published version of the manuscript.

## SUPPLEMENTAL INFORMATION

Supplemental information can be found online at XXX.

## STAR★METHODS

### KEY RESOURCES TABLE

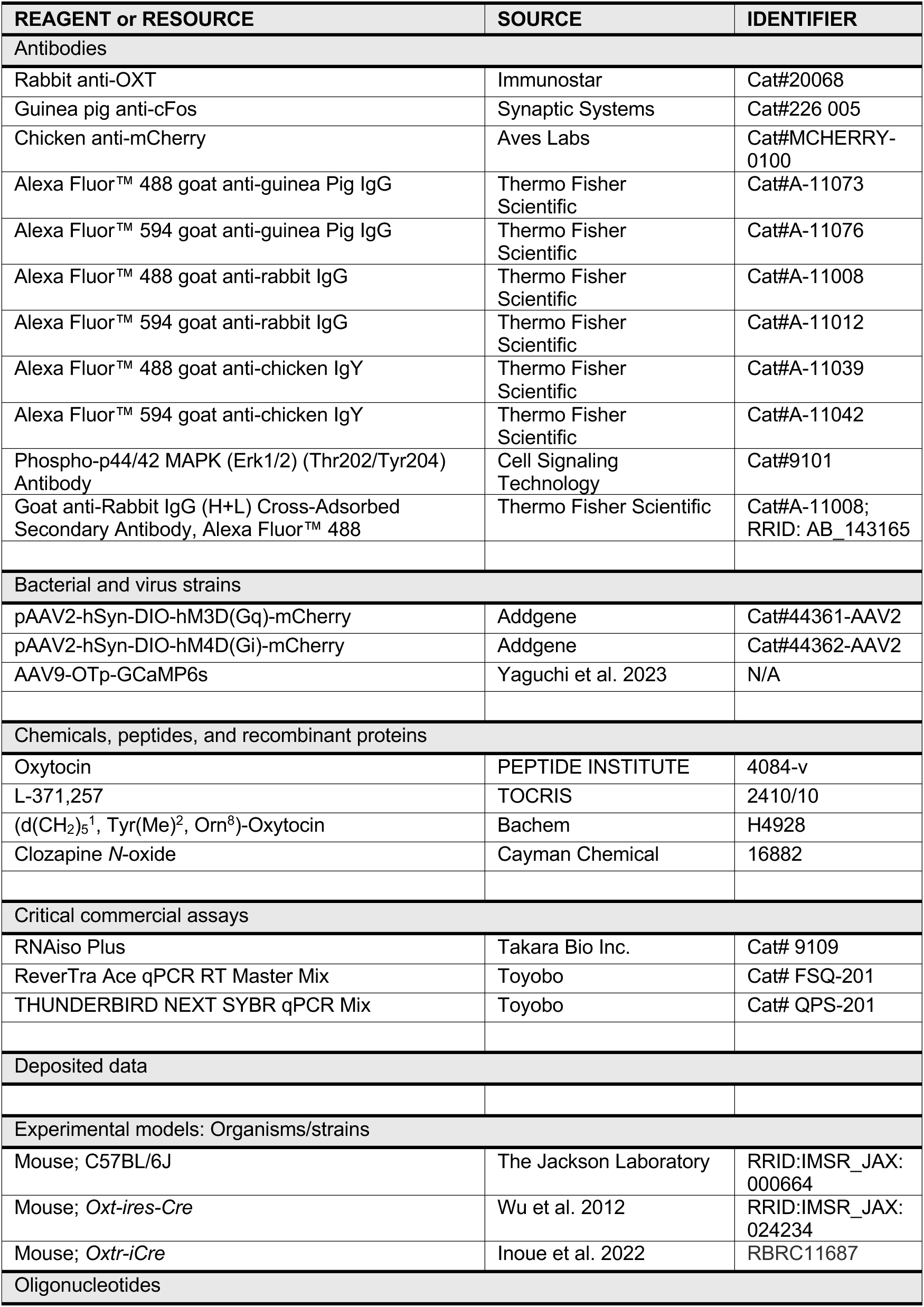
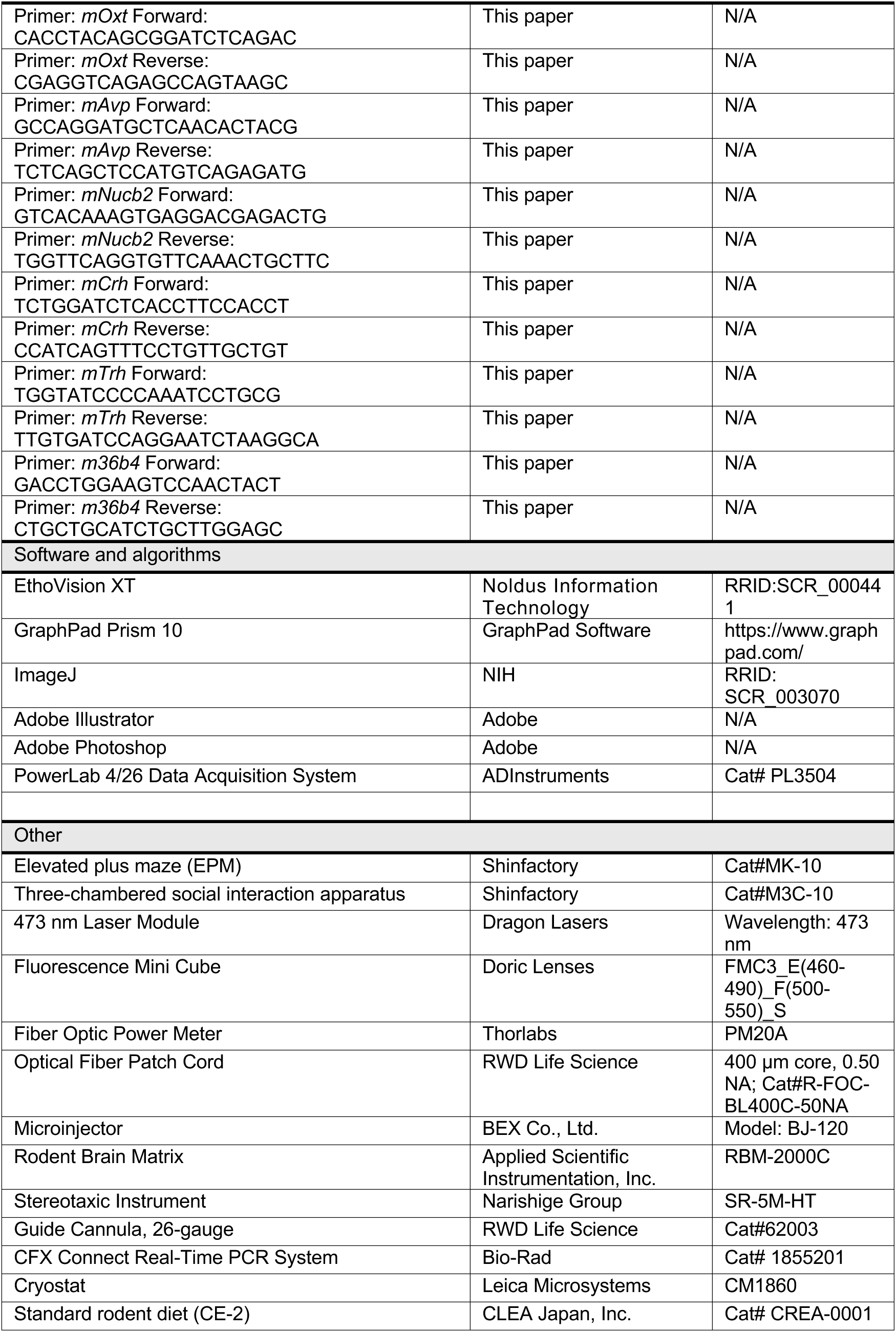

### EXPERIMENTAL MODEL AND SUBJECT DETAILS

#### Mice

Wild-type C57BL/6J male mice (The Jackson Laboratory Japan, Inc., Yokohama, Japan), *Oxt-ires-Cre* mice (a kind gift from Dr. Brad Lowell, Beth Israel Deaconess Medical Center & Harvard Medical School; ^45^, *Oxtr-iCre* mice (a kind gift from Dr. Kazunari Miyamichi, RIKEN Center for Biosystems Dynamics Research; ^46^ were obtained. *Oxt-ires-Cre* mice had a 129/BL6 mixed background, maintained with five backcrosses to C57BL/6J mice.

All mice were group-housed under controlled temperature (22.5 ± 2°C), humidity (55 ± 10%), and a 12-h light/dark cycle (lights on 07:30–19:30). Standard laboratory chow (CE-2, CLEA Japan, Tokyo, Japan) and water were available *ad libitum*. Purchased mice were allowed to acclimate to the facility for at least one week before any procedures. All mice (age 8-30 weeks) were used for all behavioral studies and were adequately habituated to handling prior to the experiments. The animal experiments were carried out after receiving approval from the Institutional Animal Experiment Committee of the Kyoto Prefectural University and in accordance with the Institutional Regulations for Animal Experiments (approval number: KPU070217-RC-3, KPU070217-RC2, KPU070217-RC1 and KPU070217-RC5).

### METHOD DETAILS

#### Surgeries

All surgical procedures were performed under anesthesia induced by IP injection of a three-drug combination (MMB anesthetic) consisting of medetomidine (0.75 mg/kg; Nippon Zenyaku Kogyo, Koriyama, Japan), midazolam (4.0 mg/kg; Maruishi Pharmaceutical, Osaka, Japan), and butorphanol (5.0 mg/kg; Meiji Seika Pharma, Tokyo, Japan). After surgery, atipamezole (0.75 mg/kg, IP; Nippon Zenyaku Kogyo) was administered to reverse anesthesia, and mice were kept on a heating pad maintained at 38°C until fully ambulatory.

#### Stereotactic surgery

Mice were anesthetized with the MMB anesthetic and head-fixed in a stereotaxic frame (Narishige, Tokyo, Japan). Either adeno-associated virus (AAV) was microinjected, or an optical fiber or stainless-steel cannula was implanted into the target brain region according to the mouse brain atlas of Franklin and Paxinos.

For chemogenetic experiments, AAV2-hSyn-DIO-hM4Di-mCherry (1.0×10^12^ vg/mL) was bilaterally injected into the paraventricular nucleus of the hypothalamus (PVH; 800 nl per site) of *Oxt-ires-Cre* male mice using a microinjector (BJ-120; BEX, Tokyo, Japan). For fiber photometry experiments, AAV9-OTp-GCaMP6s (2.91×10^12^ vg/ml, 400 nl; ^22^ was unilaterally injected into the PVH of C57BL/6J male. The coordinates for the PVH were (in mm from bregma): AP –0.75, ML ± 0.25, DV –4.75. For implantation of optical fibers or stainless-steel cannulas, an optical fiber (400 µm core, 0.50 NA; RWD Life Science, Shenzhen, China) was implanted directly above the PVH injection site (DV –4.65) immediately following AAV9-OTp-GCaMP6s delivery for fiber photometry. For intracerebroventricular (ICV) drug administration, a stainless-steel guide cannula (26-gauge; RWD Life Science, Shenzhen, China) was unilaterally implanted into the right lateral ventricle of C57BL/6J male mice at the following coordinates (in mm from bregma): AP –0.25, ML 1.00, DV –2.50. All implants were secured to the skull using dental cement. Mice were tested after at least 3 weeks of recovery from AAV injection and 1 week after ICV cannula implantation.

#### Subdiaphragmatic vagotomy

Subdiaphragmatic vagotomy was performed as previously described ^9^. In brief, under MMB anesthesia, a midline abdominal incision was made in C57BL/6J male mice to expose the subdiaphragmatic vagal nerve trunks running along the esophagus. The ventral, dorsal, or both subdiaphragmatic trunks of the vagus nerve were exposed and transected. In the sham operation group, these vagal trunks were exposed but not cut. After bilateral subdiaphragmatic vagotomy, mice were maintained on a nutritionally complete liquid diet (Ensure-H; Abbott Japan, Tokyo, Japan), whereas after unilateral vagotomy, mice were maintained on standard laboratory chow. One to two weeks post-operation, animals underwent behavioral testing and brain gene expression analysis.

#### Microinjection of AAV vector into the nodose ganglion

Microinjection of AAV vector into the nodose ganglion (NG) was performed with modifications to previously described methods ^16,47^. *Oxtr-iCre* mice were anesthetized with the MMB anesthetic, and the left or right NG was surgically exposed. AAV2-hSyn-DIO-hM3Dq-mCherry (1.0 ∼ 3.0 × 10^12^ vg/ml, 1 µl per site, containing 0.05% Fast Green for enhance visualization) was injected using a microinjector (BJ-120; BEX). Behavioral experiments were conducted at least 3 weeks post-surgery. Following the completion of behavioral testing, immunohistochemistry was performed to confirm hM3Dq-mCherry expression.

#### Preparation of reagents

Oxytocin (Oxt) and the Oxt receptor antagonist [d(CH_2_)_5_^1^,Tyr(Me)^2^,Orn^8^]-oxytocin (H4928); OVT were dissolved directly in saline. The oxytocin receptor antagonist L-371,257, which has low blood–brain barrier permeability, and the DREADD ligands clozapine *N*-oxide (CNO) were first dissolved in DMSO and then diluted with saline. The final concentrations of DMSO in the injection solutions were 20% for L-371,257 and 1% for CNO.

#### Elevated plus maze (EPM)

The EPM (MK-10; Shinfactory, Fukuoka, Japan) consisted of two opposing open arms (30 × 6 cm) and two opposing closed arms (30 × 6 cm) with 20 cm high gray polycarbonate walls, connected to a central zone (6 × 6 cm) forming a cross, and was elevated 50 cm above the floor with gray floors throughout. The gray polycarbonate material was optimal as it minimized light reflection in the video recordings. This EPM test was conducted in an independent, quiet room with controlled temperature and humidity. The tests were performed during the light phase (13:00–18:00). Mice were acclimated to this environment for at least 90 min before testing. The illumination of the experimental room was adjusted using white LED lights so that the central zone of the EPM was 50 lux.

Oxt (400 nmol/kg), L-371,257 (10 mg/kg), Oxt + L-371,257 (400 nmol/kg and 10 mg/kg), or vehicle (saline or saline containing 20% DMSO, 10 ml/kg) was administered intraperitoneally 1 h before the start of the EPM test. OVT (3.7 nmol in 2 µl) or saline was administered intracerebroventricularly 75 min before the test. CNO (1 mg/kg) or vehicle (1% DMSO in saline) was administered intraperitoneally 2 h before the EPM test for hM4Di or 1 h before the test for hM3Dq. After drug administration, mice were placed individually in the center of the maze and allowed to freely explore. Mouse behavior was recorded for 10 min using an area scan camera (107653, Basler, Ahrensburg, Germany) equipped with a wide-angle lens (2000034831, Basler) covering the entire maze. Behavioral parameters, including the percentage of time spent in the open and closed arms and the total distance traveled, were recorded and analyzed using EthoVision XT 16 (Noldus Information Technology, Wageningen, the Netherlands). After each session, the maze was wiped with 50 ppm sodium hypochlorite solution (super hypochlorous water), followed by 70% ethanol, and then air-dried for 5 min before testing the next mouse.

#### Three-chambered social interaction test (SIT)

The three-chambered social interaction apparatus consisted of a rectangular box (internal dimensions: W60 × D40 × H23 cm) divided into three equal compartments by transparent acrylic walls with openings that allowed free movement between chambers (Figure S1A; M3C-10, Shinfactory). Two custom-designed L-shaped wire cages (Figure S1B and S1C; gray; 10 cm per side × 20 cm height) were placed in each of the two side chambers. The L-shape target cage was used instead of conventional cylindrical or square cages to ensure that the subject mice’s nose remained visible without shadow to the overhead video camera, thereby allowing precise tracking of sniffing behavior (Figure S1D). The sniffing zone was defined as the area within 2 cm surrounding each wire target cage (Figure S1E).

Tests were conducted during the light phase (13:00–18:00). Mice were transferred to the same experimental room as used for the EPM at least 90 min before testing. The illumination was adjusted so that the center chamber was 50 lux. Test compounds were administered under the same conditions as in the EPM. Each SIT consisted of two sequential 10-min sessions separated by a 2.5-min interval: a habituation session and a social interaction session. In the habituation session, the subject mouse was placed in the central chamber and allowed to freely explore all three chambers for 10 min, with both target cages kept empty. After this session, the subject mouse was placed in a clean holding cage for a 2.5-min break, during which the apparatus and cages were wiped with 50 ppm sodium hypochlorite solution and 70% ethanol. In the next social interaction session, a novel sex-matched conspecific mouse was placed in the left-side target cage, while the right-side cage remained empty, and the subject mouse was again allowed to explore freely for 10 min. Behavioral parameters, including the time the subject mouse’s nose remained within each sniffing zone and the total distance traveled, were recorded and analyzed using the same video tracking system and EthoVision XT 16 as described for the EPM. The time spent in the sniffing zone of the target mouse was used as the primary index of social interaction.

#### Measurements of food intake

Mice were individually housed and habituated for at least one week to a powdered CE-2 standard chow diet (CLEA Japan) in a feeding box (Shinano Manufacturing Co., Ltd., Tokyo, Japan), and to gentle handling. Food was removed at 16:30 with free access to water (3 h fasting), and feeding experiments were conducted starting at 19:30. In intact, sham-operated, and vagotomized mice, Oxt (400 nmol/kg), saline (10 ml/kg), L-371,257 (10 mg/kg), or a combination of Oxt and L-371,257 was administered intraperitoneally at 19:20. In *Oxt-ires-Cre* mice expressing hM4Di in the PVH, CNO (3 mg/kg) or vehicle containing 1% DMSO (10 ml/kg) was administered intraperitoneally 1 h prior to Oxt or saline injection. In *Oxt-ires-Cre* mice expressing hM3Dq in the NG, CNO (1 mg/kg) or vehicle was administered intraperitoneally at 19:20. The feeding box, including the powdered food and food spillage, was weighed 1 to 14 h after the start of refeeding. The cumulative energy intake at each time point was calculated based on the weight of consumed CE-2 chow (3.4 kcal/g) and expressed as kcal.

#### Fiber photometry

Fiber photometry recordings were conducted at least 3 weeks after AAV injection. On the day of the experiments, mice were fasted for at least 3 h before testing, and recordings were performed between 10:30 and 16:30. After a 15-min baseline recording, all mice received an IP injection of saline (10 ml/kg), Oxt (400 nmol/kg), Oxt + L-371,257 (400 nmol/kg and 10 mg/kg), or Oxt solution containing 20% DMSO as the control for Oxt + L-371,257. Excitation light (473 nm; Dragon Lasers, Changchun, China) was delivered through the implanted optical fiber to record Ca^2+^ signals from PVH^Oxt^ neurons. The resulting GCaMP6s fluorescence was collected and filtered (500–550 nm) using an integrated fluorescence mini cube (Doric Lenses, Quebec, QC, Canada). The laser power at the fiber tip was adjusted to approximately 20 µW. Signals were acquired at 1 kHz, passed through a 20 Hz low-pass filter, and downsampled to 5 Hz for analysis using a PowerLab data acquisition system (4sp; ADInstruments, Bella Vista, NSW, Australia).

The change in cytosolic Ca^2+^ concentration from baseline (ΔF/F) was calculated as ΔF/F(%)=100×(F_t_−F_baseline_)/F_baseline_, where F_t_ is the fluorescence at a given time point t, and F_baseline_ is the average fluorescence during a 15-min baseline period immediately preceding the stimulus. To visualize the activity profile around an injection event, peri-event traces were generated by extracting ΔF/F data from –15 min to 60 min relative to the injection. These data were then averaged into 3 min bins for plotting. The IAUC as calculated as the cumulative sum of the binned ΔF/F values from 0 to 30 min post-injection, with values below baseline treated as negative components.

#### Quantitative PCR analysis

Mice were fasted from 9:00 on the day of the experiment, and Oxt (400 nmol/kg) or saline was IP administered at 12:00. At 14:00, mice were anesthetized with 30% isoflurane and subsequently decapitated. Following rapid removal, brains were placed in an ice-cold brain matrix (Applied Scientific Instrumentation, Inc., Eugene, OR, USA). A coronal slice (Bregma −0.40 to −1.4 mm) was obtained using a razor blade, and the PVH and supraoptic nucleus (SON) were subsequently microdissected under microscopic guidance and immediately immersed in RNAlater solution (Sigma). Total RNA was isolated from collected tissues using RNAiso Plus (Takara Bio Inc., Shiga, Japan) according to the manufacturer’s protocol. Subsequently, cDNA was synthesized from DNase-treated total RNA using the ReverTra Ace qPCR RT Master Mix (Toyobo, Osaka, Japan). Real-time PCR was performed using the THUNDERBIRD NEXT SYBR qPCR Mix (Toyobo) on a CFX Connect Real-Time PCR system (Bio-Rad, Hercules, CA, USA). The cycle threshold (Ct) value for each reaction was determined by the system software. Gene expression levels were calculated by first normalizing the Ct value of each target gene to that of the housekeeping gene *36b4*, yielding the ΔCt value (ΔCt = Ct_target_−Ct_36b4_). For *Oxt, Avp*, *Nucb2*, *Crh* or *Trh* expression of feeding-related genes in the PVH and SON, ΔCt values were directly compared between experimental groups. For analyzing the effect of Oxt treatment on *Oxt*, *Avp (arginine vasopressin)*, *Nucb2 (nucleobindein-2)*, *Crh (corticotropin-releasing hormone)* or *Trh (thyrotropin-releasing hormone)* expression, relative expression levels were calculated using the comparative Ct (ΔΔCt) method, where results were expressed as fold-change relative to the corresponding saline-injected control group.

#### Immunohistochemical analysis

Immunohistochemistry for phosphorylated ERK1/2 (pERK1/2) in the NG and NTS was performed as described below. Oxt (400 nmol/kg), L-371,257 (10 mg/kg), a combination of Oxt and L-371,257, or saline containing 20% DMSO was IP injected at 9:00 into C57BL/6J mice that had been fasted overnight (16 h). 15 min after injection, mice were transcardially perfused with Zamboni’s solution (4% paraformaldehyde and 0.2% picric acid in 0.1 M phosphate buffer, pH 7.4) under MMB anesthesia. For immunostaining of c-Fos in PVH^Oxt^ neurons, CNO (1 mg/kg) was IP injected at 11:00 into *Oxt-ires-Cre* mice expressing hM4Di in PVH^Oxt^ neurons or into C57BL/6J mice lacking hM4Di expression, after food removal at 9:00. Oxt (400 nmol/kg) or saline was then administered IP 1 h after CNO injection. 90 min later, mice were transcardially perfused with Zamboni’s solution for fixation. This fasting and experimental schedule was aligned with that used for the EPM and SIT experiments.

Isolated NGs and brains were postfixed in the same fixative for 2 and 4 h at 4 °C, respectively, and incubated in phosphate buffer containing 30% sucrose for 48 h at 4 °C. Longitudinal sections (10 µm) of NGs were cut at 60 µm intervals and Coronal sections (40 µm) of the brain were cut at 120 µm intervals using a precision cryostat (Leica Microsystems, Wetzlar, Germany). After blocking with PBS containing 2% bovine serum albumin and 2% normal goat serum, sections were incubated overnight at 4°C with the following primary antibodies: rabbit anti–phospho-p44/42 MAPK (1:500), guinea pig anti–c-Fos (1:1000), rabbit anti–oxytocin (1:2000), and chicken anti–mCherry (1:300). Subsequently, the sections were incubated with the following secondary antibodies (1:500) for 30 min at room temperature: goat anti-guinea pig Alexa Fluor 488/594, goat anti-rabbit Alexa Fluor 488/594, and goat anti-chicken Alexa Fluor 488. Finally, sections were washed extensively in PBS and coverslipped using Fluoromount (#S3023; Dako, Carpinteria, CA, USA).

Fluorescence images were acquired using a DP75 digital camera controlled by cellSens V4.3 imaging software (Evident, Tokyo, Japan). Images were subsequently processed for brightness and contrast under identical conditions using Adobe Photoshop (Adobe, CA, USA). The number of pERK1/2-immunoreactive (IR) neurons in the NG was manually counted and averaged across four sections per mouse. Similarly, pERK1/2-IR neurons in the medial NTS (bregma –7.32 to –7.76 mm) were counted and averaged per section. In the PVH (bregma –0.58 to –1.06 mm), the numbers of c-Fos-IR and OXT-IR neurons, as well as their overlap, were counted.

Because a validated anti-oxytocin antibody was available, we examined the expression efficiency of the viral vectors in PVH^Oxt^ neurons after microinjection of AAV2-hSyn-DIO-hM4Di-mCherry or AAV9-OTp-GCaMP6s into the PVH. 62 ± 7% (n = 4) of PVH^Oxt^ neurons expressed the target protein following AAV2-hSyn-DIO-hM4Di-mCherry injection (*Oxt-ires-Cre mice*), and 77 ± 4% (n = 3) following AAV9-OTp-GCaMP6s injection (C57BL/6J mice).

### QUANTIFICATION AND STATISTICAL ANALYSIS

All data are shown as means ± SEM. Statistical analysis was performed using a two-tailed unpaired t-test, one-way ANOVA, or two-way ANOVA as appropriate. When ANOVA revealed significant main effects or interactions, group differences were assessed using Dunnett’s or Tukey’s post hoc tests. All analyses were conducted using Prism 10 (GraphPad Software, San Diego, CA, USA), and p < 0.05 was considered.

## Supplemental figure legends

**Figure S1.**
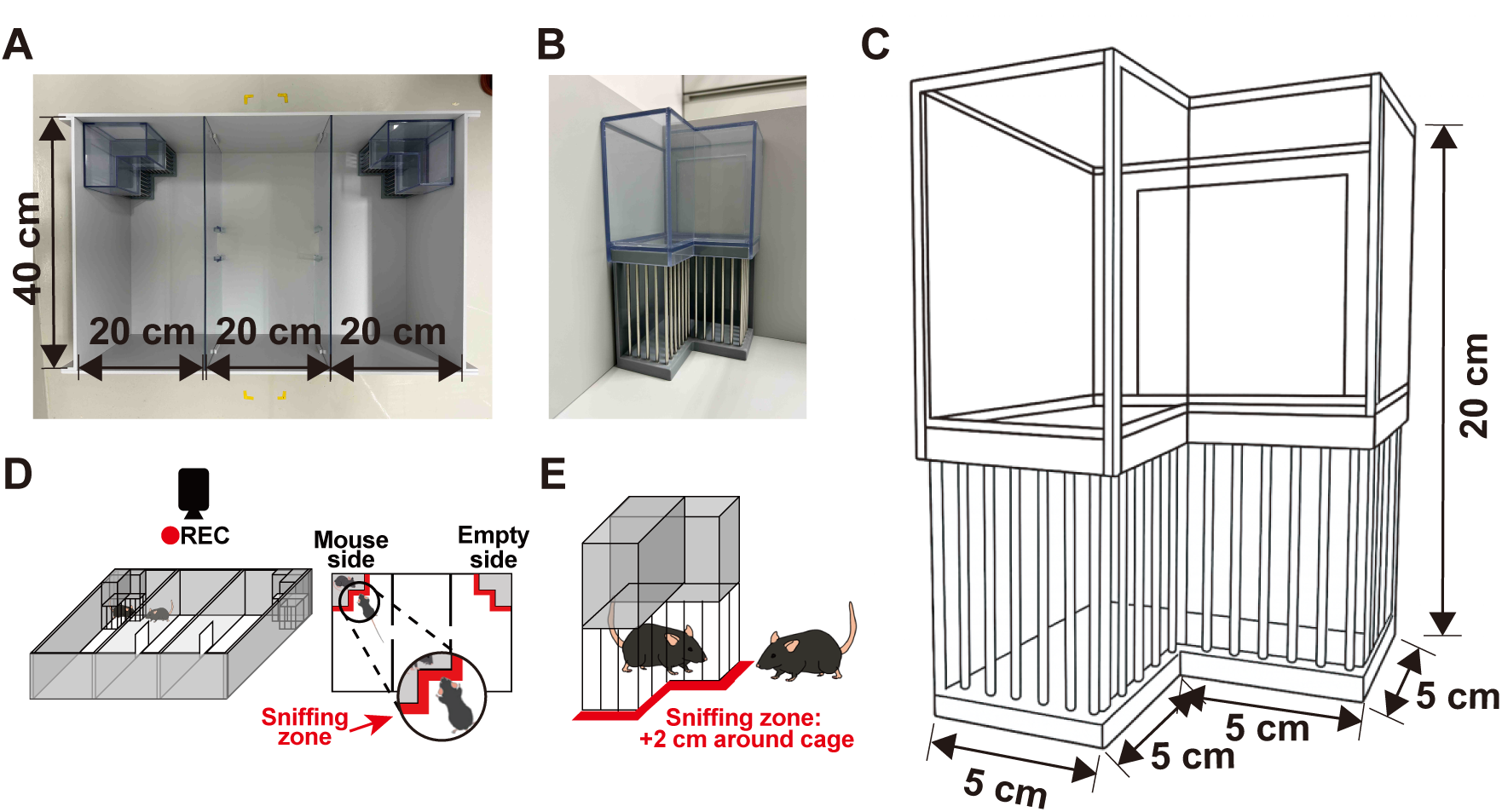
Apparatus and experimental scheme of the three-chamber social interaction test (SIT). To improve the accuracy of behavioral assessment, we used a modified system in which a specially designed L-shaped wire cage was placed in the corner of the chamber. This configuration allowed reliable detection of sniffing behavior by video recording while minimizing shadow artifacts. **(A)** Overhead view of the SIT apparatus (internal dimensions: 20 × 40 × 23 cm × 3 chambers). **(B)** Photograph of the L-shaped wire cage used in the test. **(C)** Schematic drawing of the wire cage dimensions. To prevent the mice from climbing onto the top of the wire cage (external dimensions: 10 cm high), the upper part of the cage was covered with a polyvinyl chloride box (external dimensions: 10 cm high). **(D)** Experimental scheme of the SIT. A subject mouse was placed in the central chamber at the beginning of the test. The three chambers were connected by windows (5 cm wide × 8 cm high) located on both sides of the central chamber, allowing the mouse to move freely among the three chambers. The subject mouse was allowed to explore either the target cage (mouse side) or the empty cage (empty side). Sniffing behavior toward each cage was recorded. **(E)** Definition of the sniffing zone. The sniffing zone was defined as the area extending 2 cm around the target cage, and the time the mouse’s nose remained within this zone was measured using a video tracking system.

**Figure S2.**
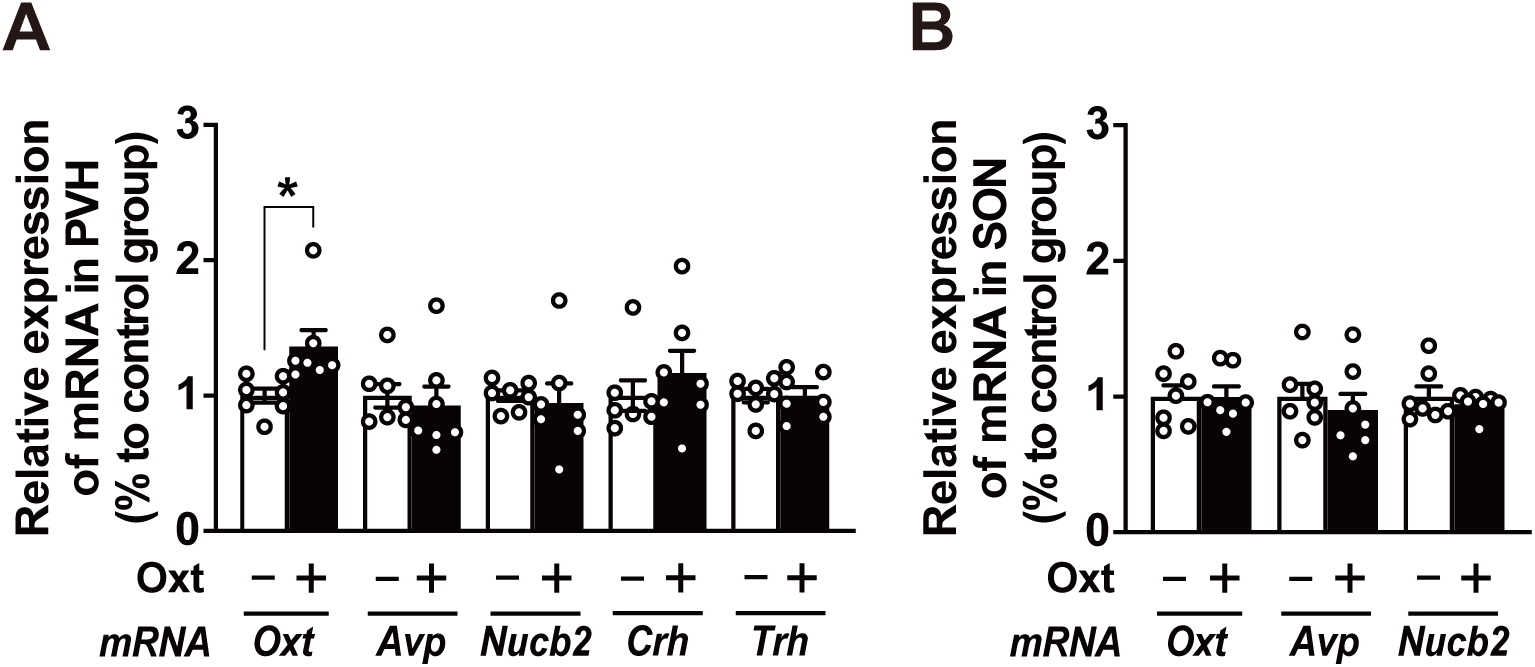
IP injection of Oxt selectively increases *Oxt* mRNA expression in the PVH but not in the SON. (**A–B**) Relative mRNA expression of neuropeptides involved in feeding regulation in PVH (**A**) or SON (**B**) 2 h after IP injection of Oxt at 400 nmol/kg or saline in C57BL6/J male mice. *Oxt*, oxytocin; *Avp*; arginine vasopressin; *Nucb2*, nucleobindein-2; *Crh*, corticotropin-releasing hormone; *Trh*, thyrotropin-releasing hormone. 36b4 was used as an internal control. n = 7. *p < 0.05 by unpaired t-test (A).

